# Protective Role of Galectin-3 in Prion Disease Through Regulation of Microglial PrP^Sc^ Uptake

**DOI:** 10.64898/2026.06.11.731738

**Authors:** Natallia Makarava, Tarek Safadi, Narayan P. Pandit, Olga Mychko, Olga Bocharova, Kara Molesworth, Marta M. Lipinski, Ilia V. Baskakov

## Abstract

Microglia constitute a major innate defense mechanism against prion infection; however, the molecular pathways regulating microglial responses during disease progression remain incompletely understood. Galectin-3 (Gal3), a β-galactoside-binding lectin associated with reactive microglia in multiple neurodegenerative disorders, has been implicated in phagocytosis, inflammatory signaling, and lysosomal homeostasis. Here, we investigated the role of Gal3 in prion disease pathogenesis using prion-infected mice. Gal3 expression was undetectable in healthy brain but became upregulated beginning at late preclinical stages, increasing with disease progression. Gal3 localized predominantly to a subpopulation of reactive IBA1-positive microglia, particularly within the thalamus, and inversely correlated with expression of the homeostatic microglial markers P2Y12 and TMEM119, consistent with acquisition of a reactive phenotype. Microglia engaged in neuronal envelopment displayed elevated Gal3 expression during terminal disease. Constitutive deletion of Gal3 significantly accelerated clinical disease progression without altering total PrP^Sc^ accumulation, reactive gliosis, neuronal envelopment, or overall microglial and astrocytic activation. However, Gal3 deficiency markedly reduced microglial uptake of PrP^Sc^, resulting in a lower intracellular-to-extracellular PrP^Sc^ ratio, supporting a role for Gal3 in phagocytic sequestration of prions. In contrast, Gal3 deficiency did not impair lysosomal activity, lysosomal membrane integrity, or expression of genes involved in lysosomal repair pathways. Likewise, selective inhibition of autophagy in myeloid cells exerted only minor effects on disease progression. Collectively, these findings identify Gal3 as a sensitive marker of reactive microglia that contributes to microglial uptake of PrP^Sc^ and exerts a protective role during prion disease progression.

## Introduction

Prion diseases, also known as transmissible spongiform encephalopathies (TSEs), comprise a group of fatal transmissible neurodegenerative disorders affecting both humans and animals [1]. Despite extensive research efforts, no effective therapies are currently available. Prion diseases are caused by prions, or PrP^Sc^, the misfolded and aggregated form of the cellular prion protein (PrP^C^) a host-encoded sialoglycoprotein [2]. Disease progression is driven by the self-propagating conversion of PrP^C^ into β-sheet-rich PrP^Sc^ conformers, leading to the accumulation and spread of prions throughout the central nervous system (CNS) [3]. Progressive neurodegeneration in prion-infected individuals and animals is closely associated with the accumulation of PrP^Sc^ and its neurotoxic effects on neurons [4–10].

Microglia are the principal innate immune cells of the CNS, where they continuously monitor the brain microenvironment and mediate the clearance of damaged cells and protein aggregates. Accumulating evidence supports the concept that microglia constitute a major host defense mechanism against prion infection [11–13]. Indeed, depletion of microglia either before prion inoculation or during the early stages of disease accelerates disease progression and shortens survival time [14–17]. However, other studies suggest that microglia may exert deleterious effects during later disease stages. Partial inhibition of microglial proliferation and reactivity during the preclinical phase delayed the onset of neurological symptoms and prolonged survival [18]. Similarly, pharmacological suppression of microglial activation at disease onset reduced reactive gliosis and extended survival in humanized mice infected with human prions [19]. Together, these findings support the emerging view that microglia may transition from protective to pathogenic phenotypes as neurodegeneration progresses, a concept increasingly recognized across multiple neurodegenerative disorders, including Alzheimer’s disease (AD).

Galectin-3 (Gal3) is a multifunctional β-galactoside-binding lectin implicated in microglial activation, innate immune signaling, and tissue remodeling within the CNS [20–22]. Gal3 preferentially recognizes β-galactoside-containing glycans and exhibits high affinity for branched complex N-glycans lacking terminal sialylation [23–26]. Under physiological conditions, Gal3 expression in the CNS is low; however, injury, infection, and protein misfolding trigger robust upregulation of Gal3, particularly in activated microglia [20–22, 27]. Gal3 functions both intracellularly and extracellularly, acting as an opsonin. By binding to TLR4 and integrins, Gal3 amplifies pro-inflammatory signaling cascades and promotes the transition of microglia to an amoeboid, phagocytic phenotype [28]. Through its glycan-binding properties, Gal3 is believed to enhance the uptake of extracellular matrix components, apoptotic cells, myelin debris, and misfolded protein aggregates, contributing to tissue remodeling during chronic neuroinflammation.

Intracellularly, Gal3 has been implicated in the maintenance of lysosomal integrity. Upon lysosomal membrane damage, Gal3 rapidly accumulates on ruptured lysosomes, where it recruits components of the ESCRT machinery and autophagy pathways to coordinate membrane repair and lysophagy [29, 30]. These activities suggest that Gal3 may play an important role in regulating cellular responses to proteotoxic stress and impaired protein degradation.

In neurodegenerative disorders, including AD, Parkinson’s disease, Huntington’s disease, multiple sclerosis, and prion diseases, Gal3 is consistently upregulated in reactive microglia [20, 27, 31–34]. Single-cell transcriptomic studies have identified Gal3 as a characteristic marker of disease-associated microglia (DAM) [21]. However, the functional consequences of Gal3 upregulation appear to be context-dependent. While Gal3 can facilitate debris clearance and tissue repair, sustained Gal3 expression may perpetuate inflammatory signaling, promote synaptic remodeling, and contribute to chronic glial activation [22, 31, 33]. Elevated Gal3 levels in cerebrospinal fluid and brain tissue correlate with disease progression and neuroinflammatory burden, highlighting its potential utility as both a biomarker and therapeutic target [27, 35].

Consistent with these observations, genetic deletion or pharmacological inhibition of Gal3 attenuated disease phenotypes in several experimental models of neurodegeneration. In the 5XFAD mouse model of AD, Gal3 expression was markedly elevated in plaque-associated microglia, whereas Gal3 deletion reduced amyloid-β burden and improved cognitive performance [31]. Gal3 depletion also suppressed tau propagation, neuroinflammation, and associated functional impairments [33]. In models of retinal degeneration, inhibition of Gal3 reduced microglial activation and delayed neurodegeneration [36]. Similarly, in Huntington’s disease models, Gal3 deletion diminished inflammatory responses, reduced huntingtin aggregation, improved motor function, and prolonged survival [27]. Collectively, these findings suggest that Gal3 contributes to neurodegenerative pathology and may represent a promising therapeutic target for slowing disease progression.

In prion disease, Gal3 expression is strongly upregulated in affected brain regions characterized by prion deposition and neurodegeneration [34, 37–41]. Elevated Gal3 expression localizes predominantly to myeloid cells, including reactive microglia [39, 41]. Nevertheless, it remains unclear whether Gal3 serves merely as a marker of microglial activation or functions as an active disease-modifying factor capable of influencing prion pathogenesis. In the present study, we investigated the impact of Gal3 deletion on prion disease progression and examined the role of Gal3 in regulating reactive microglial responses associated with prion-induced neurodegeneration.

## Methods

### Animals

10% (w/v) SSLOW brain homogenates (BH) for inoculations were prepared in PBS, pH 7.4, using glass/Teflon homogenizers attached to a cordless 12 V compact drill. Immediately before inoculation, each inoculum was further dispersed by 30 seconds of indirect sonication at approximately 200 watts in a microplate horn of a sonicator (Qsonica, Newtown, CT) and diluted in PBS, pH 7.4. Brain-derived materials were inoculated as 1% via i.c. (intracerebral) or i.p. (intraperitoneal) route, as noted. Male and female Gal3 KO mice (#006338, Jackson Laboratory, Bar Harbor, ME) and C57BL/6J (Veterinary Resources, University of Maryland, Baltimore, Maryland, USA), 6–7 weeks old, received 200 µl of inoculum i.p. or 20 µl of inoculum i.c. under 3% isoflurane anesthesia and were regularly monitored for clinical signs, which included clasping hind legs, difficulty walking, abnormal gait, nesting problems, and weight loss. Naïve age-matched C57BL/6J and Gal3 KO mice were used as non-infected controls. The mice were considered terminal and euthanized when they were unable to rear and/or lost 20% of their weight. Becn1 cKO (Becn1^flox/flox^Lyz2^Cre/Cre^) and control (Lyz2^Cre/Cre^) female mice [42] were inoculated with 1% SSLOW ic and assessed as above. P2Y12 KO and CD11b KO mice were described previously [40, 43].

The study was carried out in strict accordance with the recommendations in the Guide for the Care and Use of Laboratory Animals of the National Institutes of Health (National Academies Press, 2011). The animal protocol was approved by the Institutional Animal Care and Use Committee of the University of Maryland, Baltimore (assurance number A32000-01; permit number 0215002).

### Sex as a biological variable

Majority of the experiments were performed on male and female mice, and similar findings are reported for both sexes. Becn1 cKO (Becn1^flox/flox^; Lyz2^Cre/Cre^) and control (Lyz2^Cre/Cre^) mice were only females.

### Antibodies

Primary antibodies used for immunofluorescence, immunohistochemistry, and immunoblotting were as follows: mouse monoclonal anti-prion protein, clone SAF-84 (189775, Cayman Chemicals, Ann Arbor, MI); rabbit monoclonal anti-prion protein, clone 3D17 (ZRB1268, Millipore Sigma, Rockville, MD), rabbit polyclonal anti-IBA1 (013-27691, Fujifilm Wako Chemicals USA, Richmond, VA); goat polyclonal anti-IBA1 (NB100-1028, Bio-techne Minneapolis, MN); rat monoclonal anti-Gal3, clone M3/38 (sc-23938, Santa Cruz Biotechnology Inc.); rabbit anti-P2Y12 (#55043A, Anaspec, Fremont, CA); TMEM119 (#90840, Cell Signaling Technology, Danvers, MA), rabbit anti-Vim (#5741, Cell Signaling Technology); mouse monoclonal anti-NeuN, clone A60 (MAB377, Millipore Sigma); rat monoclonal anti-LAMP1, clone 1D4B (121601, BioLegend, San Diego, CA); rabbit polyclonal anti-CD11b (ab128797, Abcam, Waltham, MA); rabbit polyclonal anti-IFITM3 (11714-1-AP, Proteintech, Rosemont, IL); rabbit monoclonal anti-cathepsin D (69854, Cell Signaling Technology), goat polyclonal anti-GPNMB (AF2320, Bio-techne), and mouse monoclonal anti–β-actin, clone AC-15 (A5441, Millipore Sigma);. The secondary antibodies for immunofluorescence were Alexa Fluor 488, 546, and 647 labeled (Thermo Fisher Scientific, Waltham, MA).

### Immunofluorescence and DAB staining of mouse brains

Formalin-fixed brains (sagittal or coronal 3 mm slices) were treated for 1 hour in 96% formic acid before being embedded in paraffin using standard procedures; 4 μm sections produced with Leica RM2235 microtome (Leica Biosystems, Deer Park, IL) were mounted on Superfrost Plus Microscope slides (22-037-246, Thermo Fisher Scientific) and processed for immunohistochemistry according to standard protocols. To expose epitopes, slides were subjected to 20 minutes of hydrated autoclaving at 121°C in citrate buffer, pH6.0, antigen retriever (C9999, Sigma-Aldrich, St. Louis, MO). For the detection of disease-associated PrP, an additional 3 minute treatment in concentrated formic acid was applied.

For immunofluorescence, an Autofluorescence Eliminator Reagent (2160, Sigma-Aldrich) and Signal Enhancer (11932, Cell Signaling Technology) were used on slides according to the original protocols to reduce background fluorescence. Images were collected using an inverted microscope Nikon Eclipse TE2000-U (Nikon Instruments Inc.) equipped with an illumination system X-cite 120 (EXFO Photonics Solutions Inc.) and a cooled 12-bit CoolSnap HQ CCD camera (Photometrics), or Leica Mica widefield microscope (Leica Microsystems, Boston, MA). Images were processed using ImageJ software (1.54p, NIH). Analyze Particles function of ImageJ was used for segmentation and defining regions of interest (ROIs) for the calculation of individual cell values. Cathepsin D and LAMP1 colocalization was performed with JACOP plugin of ImageJ. Full-brain tile scan images were acquired and stitched using Leica MICA widefield microscope with the 20× dry objective.

For detection with DAB, horseradish peroxidase–labeled secondary antibodies and DAB Quanto chromogen and substrate (TA125QHDX, Thermo Fisher Scientific) were used. The images were acquired with Canon G7 digital camera equipped with a microscope adaptor (Bower, NY).

### Confocal microscopy and 3D image reconstruction

Confocal images were acquired using a Leica TCS SP8 microscope using the ×40/1.30 or ×63/1.40 oil immersion objective lenses with laser lines 405, 488, 552, and 638 as needed. Image resolution was 1024 × 1024 or 2048 × 2048 pixels at a 400 Hz scan speed. For 3D reconstruction, Z-stack thickness ranged from 6.2 to 10.45 μm taken at the system-optimized number of steps. Images were processed using the LAS X, version 1.4.5.27713, and ImageJ software.

### Quantification of envelopment

To estimate the percentage of neurons enveloped by microglia and the percentage of microglial cells involved in envelopment, brains were co-immunostained for NeuN and IBA1, and immunofluorescence images from cortices of each brain were collected. Using ImageJ software, NeuN and IBA1 channel images were subjected to a threshold, converted to binary, and then individual neurons and microglia cells were identified as ROIs using the Analyze Particles function of ImageJ. The area of IBA1 signal in ROIs defined for individual neurons was measured, and the neurons were counted as undergoing envelopment if they NeuN signal overlapped by IBA1. The total number of identified NeuN+ cells was recorded as a neuronal count. The area of NeuN signal in ROIs defined for individual microglia cells was measured, and the microglia were counted as involved in envelopment if they had IBA1 signal overlapped by NeuN. The total number of identified microglia cells was recorded as a microglia count.

### Quantification of PrP^Sc^ in microglia

Brains were co-immunostained with anti-PrP (3D17 or SAF-84) and IBA1 antibodies. The images from IBA1 and anti-PrP channels were thresholded, converted to binary, and Analyze Particles function of ImageJ was used to identify individual microglia cells and PrP^Sc^ puncta on binary images from corresponding channels. The PrP^Sc^ puncta’s ROIs were measured on binary IBA1 images to classify the identified puncta as associated or not with microglia cells.

### Western blot

For Western blots, 10% (w/v) brain homogenates (BH) were prepared using RIPA Lysis Buffer (20-188, Millipore Sigma). To analyze brain-derived PrP^Sc^, BH aliquots were diluted with RIPA buffer to achieve 5% BH final concentration and treated with 20 µg/ml proteinase K (P8107S, New England BioLabs, Ipswich, MA) in the presence of 50 mM Tris, pH 7.5, and 2% Sarcosyl, for 30 min at 37 °C. To analyze other proteins, BH was diluted with RIPA buffer to 1% and proteinase digestion was omitted. The resulting samples were supplemented with 4xSDS loading buffer and heated for 10 min in a boiling water bath before loading onto NuPAGE 12% Bis-Tris gels (NP0341BOX, Thermo Fisher Scientific). Wet transfer onto PVDF membranes and probing of Western blots was done according to standard procedures. The signals were visualized by Immobilon Forte Western HRP Substrate (WBLUF0100, Millipore Sigma) or SuperSignal West pico PLUS Chemiluminescent Substrate (34577, Thermo Fisher Scientific) using Invitrogen iBright 1500 imager, and quantified with iBright Analysis software (Thermo Fisher Scientific). Intensity data were presented as normalized by actin, except for PrP^Sc^ Western blots treated with protease K.

### RT-qPCR

Total RNA was isolated from 10% BH in RIPA buffer. 100 µl BH aliquots were further homogenized within RNase-free 1.5-mL tubes in 200 µL of Trizol (15-596-026, Thermo Fisher Scientific), using RNase-free disposable pestles (12-141-364, Thermo Fisher Scientific). After homogenization, an additional 600 µL of Trizol was added to each homogenate, and the samples were centrifuged at 11,400× g for 5 min at 4 °C. The supernatant was collected, incubated for 5 min at room temperature, then supplemented with 160 µL of cold chloroform and vigorously shaken for 30 s by hand. After an additional 5-min incubation at room temperature, the samples were centrifuged at 11,400× g for 15 min at 4 °C. The top layer was transferred to new RNase-free tubes and mixed with an equal amount of 70% ethanol. Subsequent steps were performed using an Aurum Total RNA Mini Kit (7326820, Bio-Rad, Hercules, CA) following the manufacturer instructions. Isolated total RNA was subjected to DNase I digestion. RNA purity and concentrations were estimated using a NanoDrop One Spectrophotometer (Thermo Fisher Scientific). Complementary DNA (cDNA) synthesis was performed using iScript cDNA Synthesis Kit (1708890, Bio-Rad) as described elsewhere. The cDNA was amplified with CFX96 Touch Real-Time PCR Detection System (Bio-Rad) using SsoAdvanced Universal SYBR Green Supermix and the primers listed in Table 1. The PCR protocol consisted of 95 °C incubation for 2 min followed by 40 amplification cycles at 95 °C for 5 s and 60 °C for 30 s. The data were analyzed using CFX96 Touch Real-Time PCR Detection System Software (Bio-Rad).

**Table 1.**
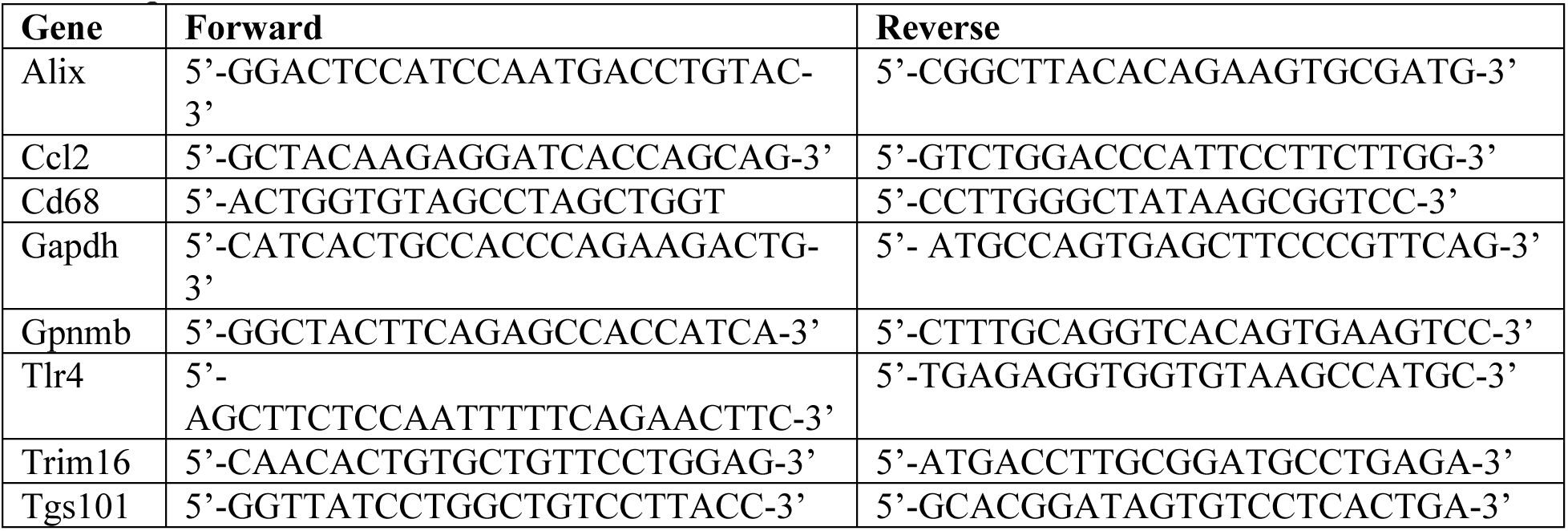
qRT-PCR Primers.

### Statistics

Statistical analyses and plotting of the data were performed using GraphPad Prism software, versions 8.4.2–10.4.1 for Windows (GraphPad Software, San Diego, CA) or Excel 2016 - version 2302. Statistical comparison of two groups with normal distribution of data points and equal variances was done with an unpaired two-tailed t-test. Alternatively, non-parametric two-tailed Mann-Whitney test was used. In case of unequal variances, the comparison of two groups with normal distribution of data points was performed with an unpaired two-tailed t-test with Welch’s correction. For statistical comparison of more than two groups, one-way ANOVAs were followed by multiple comparison tests as mentioned in Figure Legends, and the resulting p-values were reported. For statistical analysis of data presented in Superplots, means ± SD were calculated using biological replicas, where n = the number of brains or animals analyzed.

## Results

### Upregulation of Gal3 expression in a subpopulation of myeloid cells during disease progression

To establish the temporal profile of Gal3 upregulation, Gal3 expression was analyzed in C57BL/6J mice challenged intraperitoneally (i.p.) with the SSLOW prion strain. Mild upregulation was detected as early as 92 dpi; however, robust increases were observed at 122 dpi, coinciding with clinical onset [44] (Fig. 1A). Progressive elevation of Gal3 levels after 122 dpi further supported its association with disease progression. Immunohistochemical analysis revealed a regional correlation between PrP^Sc^ accumulation, microgliosis, and Gal3 immunoreactivity (Fig. 1B). Although Gal3^+^ cells were detected throughout prion-affected brain regions, Gal3 expression was especially prominent in the thalamus, the brain region most severely affected in prion-infected mice and sporadic Creutzfeldt-Jakob disease (sCJD) [34, 45–48] (Fig. 1B, Fig. S1A).

**Figure 1.**
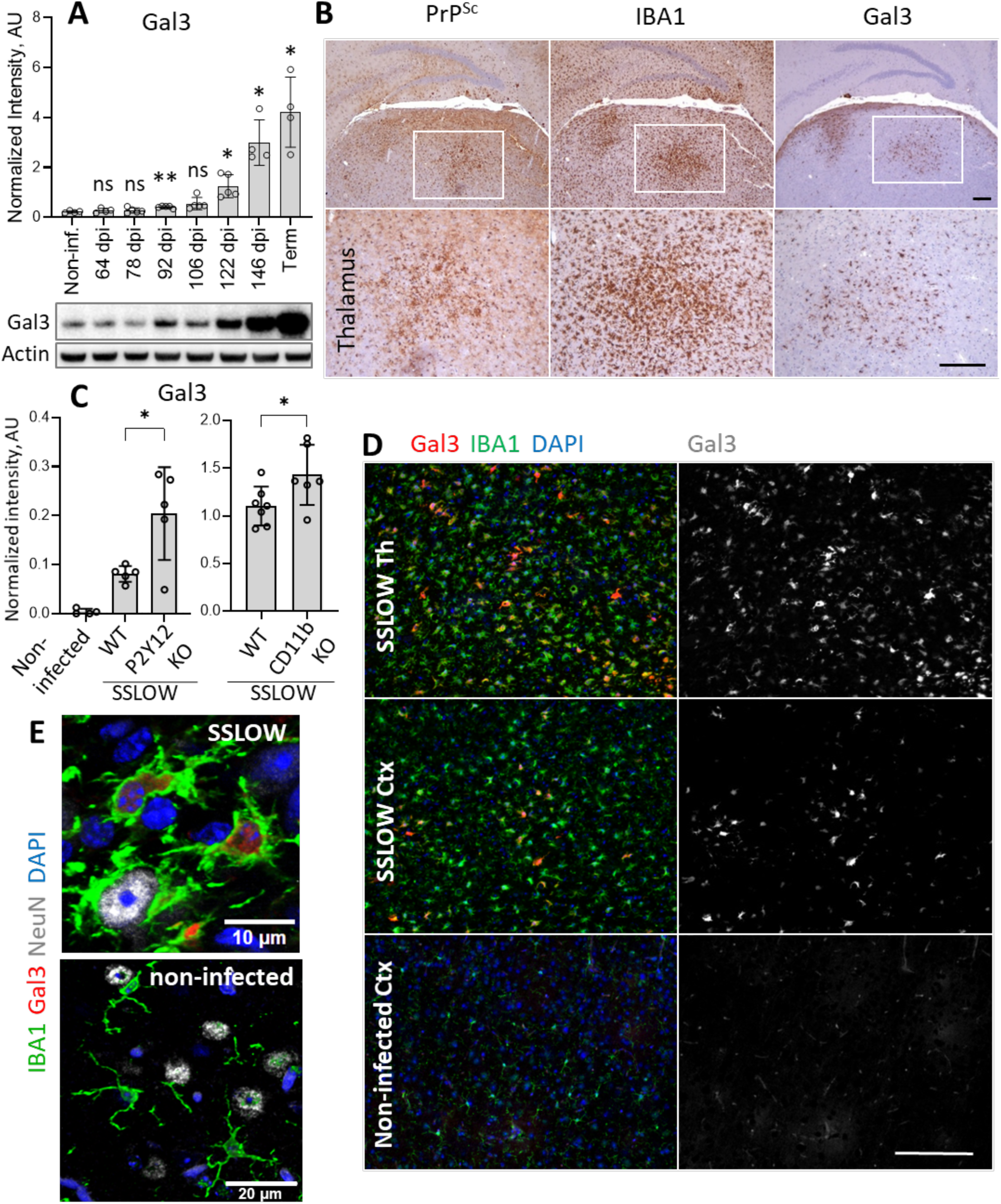
Upregulation of Gal3 expression in prion-infected mice. **A.** Upper panel: analysis of Gal3 expression in whole-brain homogenates of C57Bl/6J mice infected with SSLOW via i.p. at different stages of disease (dpi, days post inoculation; terminal state, 157-166 dpi) by Western blot. Lower panel: representative Western blots of Gal3 and actin. N=4-5 animals per time point. *p<0.05, **p<0.01, ns – not significant; comparison to non-infected brains by Brown-Forsythe and Welch’s ANOVA with Dunnett’s T3 multiple comparison test. **B.** Immunohistochemistry of C57Bl/6J mice infected with SSLOW via i.c., showing microglia reactivity (IBA1) and Gal3^+^ cells in areas with PrP^Sc^ deposition (SAF-84 antibody). Bottom row shows high-magnification views of boxed areas. Scale bar 200 µm. **C.** Western blot analysis of Gal3 expression in in whole-brain homogenates from SSLOW-infected P2Y12 knockout (KO) mice at clinical onset, CD11b KO mice at terminal stage, and corresponding WT control groups. N=5-7 animals per group, *p<0.05 by unpaired Welch’s t-test (P2Y12 KO) and Student’s t-test (CD11b KO). **D.** Immunofluorescence staining of SSLOW-infected C57Bl/6J mice illustrating Gal3 expression in a subpopulation of IBA1^+^ cells in the thalamus (Th) and cortex (Ctx), and non-infected cortex lacking detectable Gal3 signal. Scale bar 100 µm. **E.** Confocal images illustrating Gal3-positive and Gal3-negative IBA1^+^ cells in SSLOW-infected and non-infected C57Bl/6J mice, correspondingly.

Gal3 expression was highly responsive to genetic perturbations of microglia. Specifically, deletion of P2Y12, a marker of homeostatic microglia, or CD11b, a receptor involved in cell adhesion and phagocytosis, further increased Gal3 expression in prion-infected mice relative to corresponding prion-infected controls (Fig. 1C). Collectively, these data identify Gal3 as a sensitive marker of both prion infection and altered microglial phenotype in infected animals.

Co-immunostaining for IBA1 and Gal3 demonstrated that, in prion-infected brains, Gal3 was expressed by a subpopulation of IBA1^+^ myeloid cells (Fig. 1D,E). In addition, Gal3 signal co-localized with a subset of IBA1^-^/Vim^+^ cells (Fig. S1B,C), suggesting expression in reactive astrocytes under chronic inflammatory conditions. Consistent with previous observations that astrocytic responses to prions are region dependent [49–51], astrocyte-associated Gal3 expression was particularly pronounced in the striatum (Fig. S1C).

### Gal3 is upregulated in resident microglia transitioning to a reactive state

Confocal imaging of prion-infected brains revealed predominantly intracellular localization of Gal3 (Fig. 1E, Fig. 2A,C,D). To define the identity of Gal3^+^ cells, co-immunostaining for Gal3 together with TMEM119 or P2Y12, both markers of resident microglia, was performed. Under homeostatic conditions, TMEM119 and P2Y12 are highly expressed but progressively decline as microglia transition into chronically reactive states [43, 52]. In contrast, Gal3 was undetectable under homeostatic conditions (Fig. 1E, Fig. S1A) and increased progressively beginning at late preclinical stages (Fig. 1A). Consistent with these opposing expression dynamics, most Gal3^+^ cells displayed only trace levels of TMEM119 and P2Y12, whereas TMEM119^+^ or P2Y12^+^ cells that had not yet acquired a reactive phenotype exhibited little or no Gal3 signal (Fig. 2A–C). Nevertheless, IBA1^+^ cells with mild levels of both Gal3 and TMEM119/P2Y12 were also observed (Fig. 2B,C), suggesting transitional states during conversion to reactive microglia.

**Figure 2.**
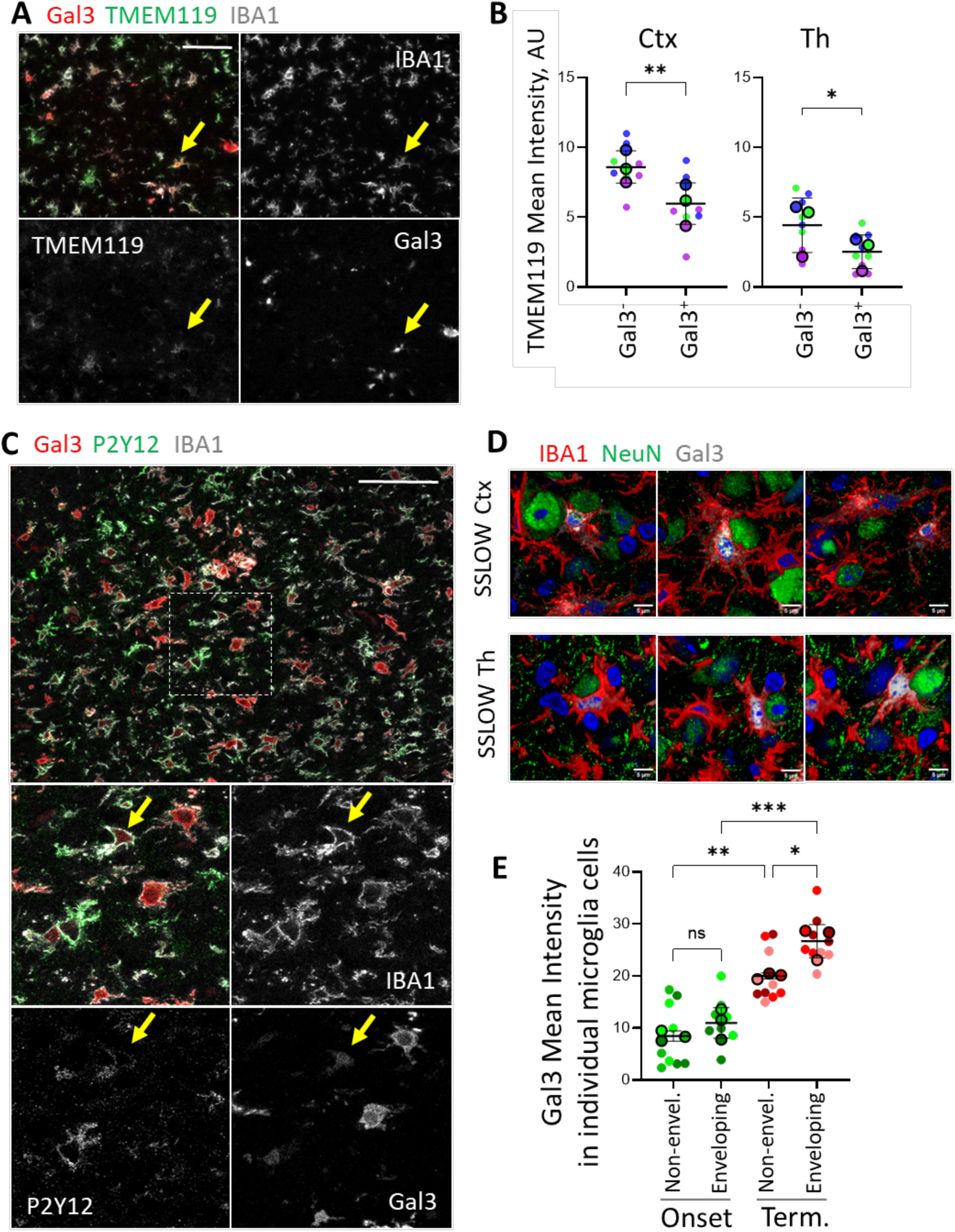
Gal3 is expressed by resident microglia including microglia that envelop neurons. **A.** Confocal images of cortex from SSLOW-infected C57Bl/6J mice immunostained for Gal3, TMEM119 and IBA1. Scale bar 50 µm. **B.** Quantification of TMEM119 mean fluorescence intensity in Gal3^-^/IBA1^+^ and Gal^+^/IBA1^+^ cells in the cortex and thalamus of SSLOW-infected C57Bl/6J mice. N=3 animals per group, *p<0.05, **p<0.01 by unpaired Student’s t-test. **C.** Confocal images of thalamus from SSLOW-infected C57Bl/6J mice stained for Gal3, P2Y12 and IBA1. Scale bar 50 µm. In **A** and **C**, arrows indicate IBA1^+^ cell co-expressing Gal3 and TMEM119, or Gal3 and P2Y12 positivity, respectively. **D.** Confocal maximum-intensity projection images of cortex (Ctx) and thalamus (Th) from SSLOW-infected mice immunostained for IBA1, NeuN and Gal3, demonstrating Gal3 expression in microglia enveloping neuronal somata. Scale bar 5 µm. **E.** Quantification of Gal3 mean fluorescence intensity in non-enveloping and enveloping IBA1^+^ microglia at clinical onset and terminal stages. N=3 brains per group *p<0.05, **p<0.01, ***p<0.001, ns – not significant by ordinary one-way ANOVA with Tukey’s multiple comparison tests. In **B** and **E**, colors represent individual brains; dots represent individual fields of view, average values for each brain are shown as circles; means ± SD are marked by black lines.

### Microglia that envelop neurons express Gal3

We previously demonstrated that, in prion disease, reactive microglia establish close body-to-body interactions with neurons, partially wrapping around neuronal somata in a process termed neuronal envelopment [40]. These interactions are dynamic, persist from minutes to hours [53], and occur across all brain regions affected by prions [40]. Neuronal envelopment is observed across multiple prion strains (SSLOW, 22L, RML, and ME7) as well as in human sCJD, suggesting a conserved mechanism [40]. Importantly, the prevalence of neuronal envelopment correlates with the severity of neuroinflammation, and genetic manipulations that increase neuronal envelopment accelerate disease progression [43]. Because Gal3 functions as an opsonin implicated in interactions between reactive microglia and neurons [54–57], we next investigated whether microglia involved in neuronal envelopment express Gal3.

Microglia engaged in neuronal envelopment exhibited a broad range of Gal3 expression, from undetectable to strongly positive (Fig. 2D, Fig. S2). This variability suggests that Gal3 is not required for initial docking of microglia to neurons, but instead may be induced following microglia–neuron contact. At clinical onset, no differences in Gal3 expression were detected between enveloping and non-enveloping microglia (Fig. 2E), potentially reflecting relatively low overall Gal3 levels at this stage. However, at terminal disease stages, microglia engaged in body-to-body interactions with neurons displayed significantly higher Gal3 levels than non-enveloping microglia (Fig. 2E). These observations prompted us to examine whether loss of Gal3 alters disease progression.

### Gal3 deficiency accelerates prion disease progression

The impact of Gal3 deficiency was investigated using constitutive Gal3 knockout (Gal3 KO) mice challenged intracerebrally (i.c.) with the mouse-adapted SSLOW strain. Clinical onset, assessed using a composite neurological score evaluating posture, mobility, clasping, and gait abnormalities, occurred at similar times in SSLOW-infected Gal3 KO and WT mice. However, following onset, disease progression was significantly accelerated in Gal3 KO animals relative to WT controls (Fig. 3A).

**Figure 3.**
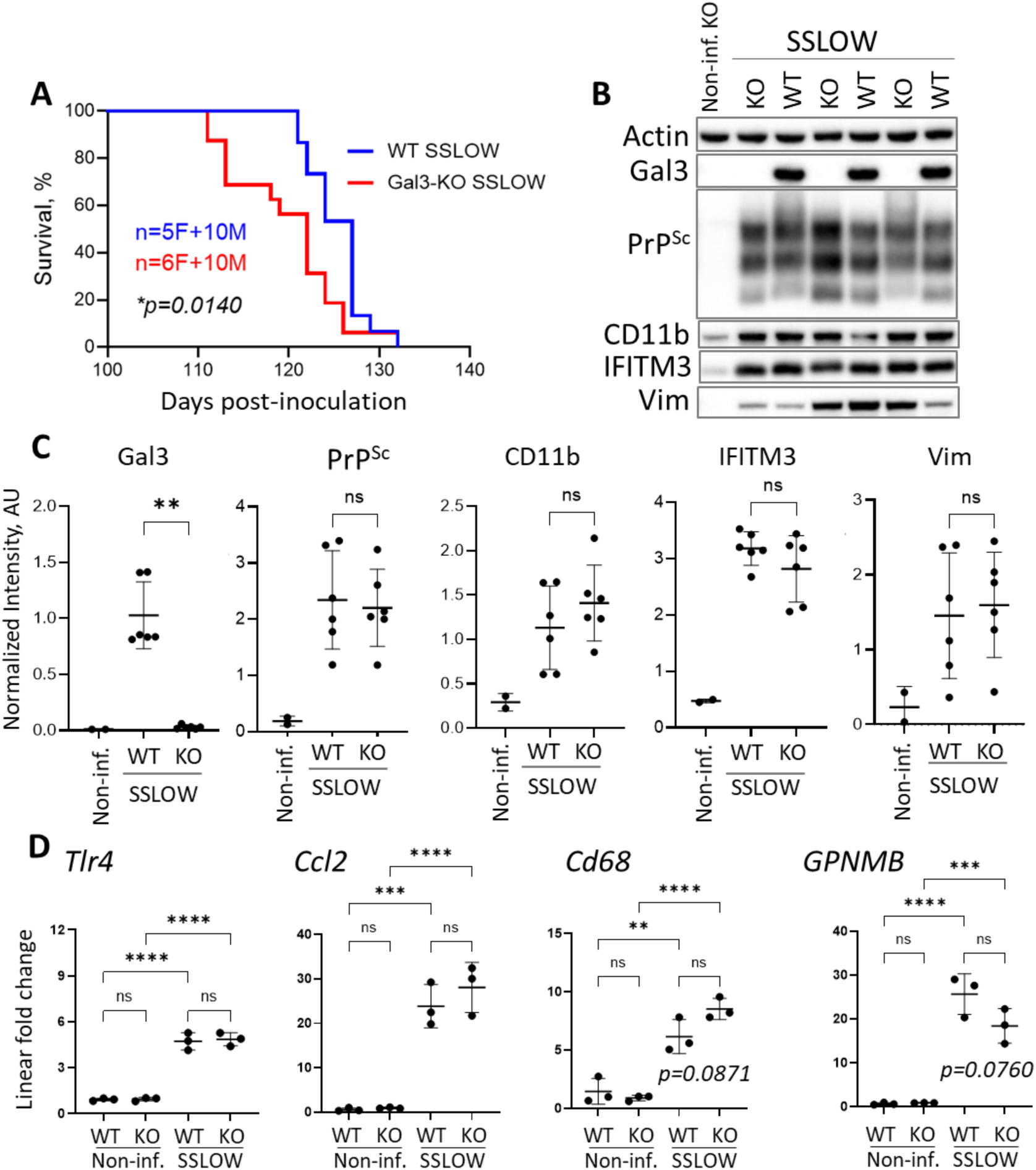
Gal3 deficiency accelerates disease progression in SSLOW-infected mice. **A.** Survival curves of Gal3 KO and WT mice inoculated with SSLOW via i.c. route. Statistical analysis was performed using the Mantel-Cox log-rank test. **B.** Representative Western blots of whole-brain homogenates from SSLOW-infected terminally ill Gal3 KO and WT control mice. **C.** Quantification of Gal3, PrP^Sc^, CD11b, IFITM3, and Vim in brains of SSLOW-infected terminally ill Gal3 KO and WT mice by Western blots. Protein levels were normalized to actin. N=6 animals per group, **p<0.01, ns – non-significant, by unpaired Student’s t-test, except for Gal3, where the non-parametric Mann-Whitney test was used. Data for non-infected brains are shown for reference (N=2). **D.** RT-qPCR analysis of gene expression in SSLOW-infected terminally ill Gal3 KO and WT mice and non-infected age-matched control groups. N=3 animals per group, **p<0.01, ***p<0.001, ****p<0.0001, ns – non-significant, by ordinary one-way ANOVA with Tukey’s multiple comparison tests.

No differences in total PrP^Sc^ accumulation were observed between Gal3 KO and WT mice at either early clinical or terminal stages (Fig. 3B,C; Fig. S3A,B). Likewise, expression of interferon-induced transmembrane protein 3 (IFITM3) and CD11b, markers upregulated in reactive microglia [58], did not differ between genotypes by Western blot analysis (Fig. 3B,C). Similarly, RT-qPCR analysis demonstrated significant upregulation of neuroinflammation-associated genes (*Tlr4, Ccl2, CD68, GPNMB*) in prion-infected relative to noninfected mice, but no significant differences between Gal3 KO and WT animals (Fig. 3D). Astrocyte reactivity, assessed by Vim expression, was also unchanged (Fig. 3B,C). Analysis of CD11b and Vim expression during early clinical stage similarly revealed no genotype-dependent differences, despite significant increases relative to noninfected mice (Fig. S3B).

### Gal3 deficiency does not alter neuronal envelopment by microglia

To investigate mechanisms underlying accelerated disease progression in Gal3 KO mice, we next examined neuronal envelopment. Neuronal and microglial densities did not differ between SSLOW-infected Gal3 KO and WT mice (Fig. 4B,D). Confocal imaging confirmed extensive neuronal envelopment in both genotypes (Fig. 4A). Quantification demonstrated that the percentage of neurons exhibiting body-to-body contacts with microglia, as well as the percentage of microglia engaged in envelopment, was comparable between Gal3 KO and WT groups (Fig. 4C,E).

**Figure 4.**
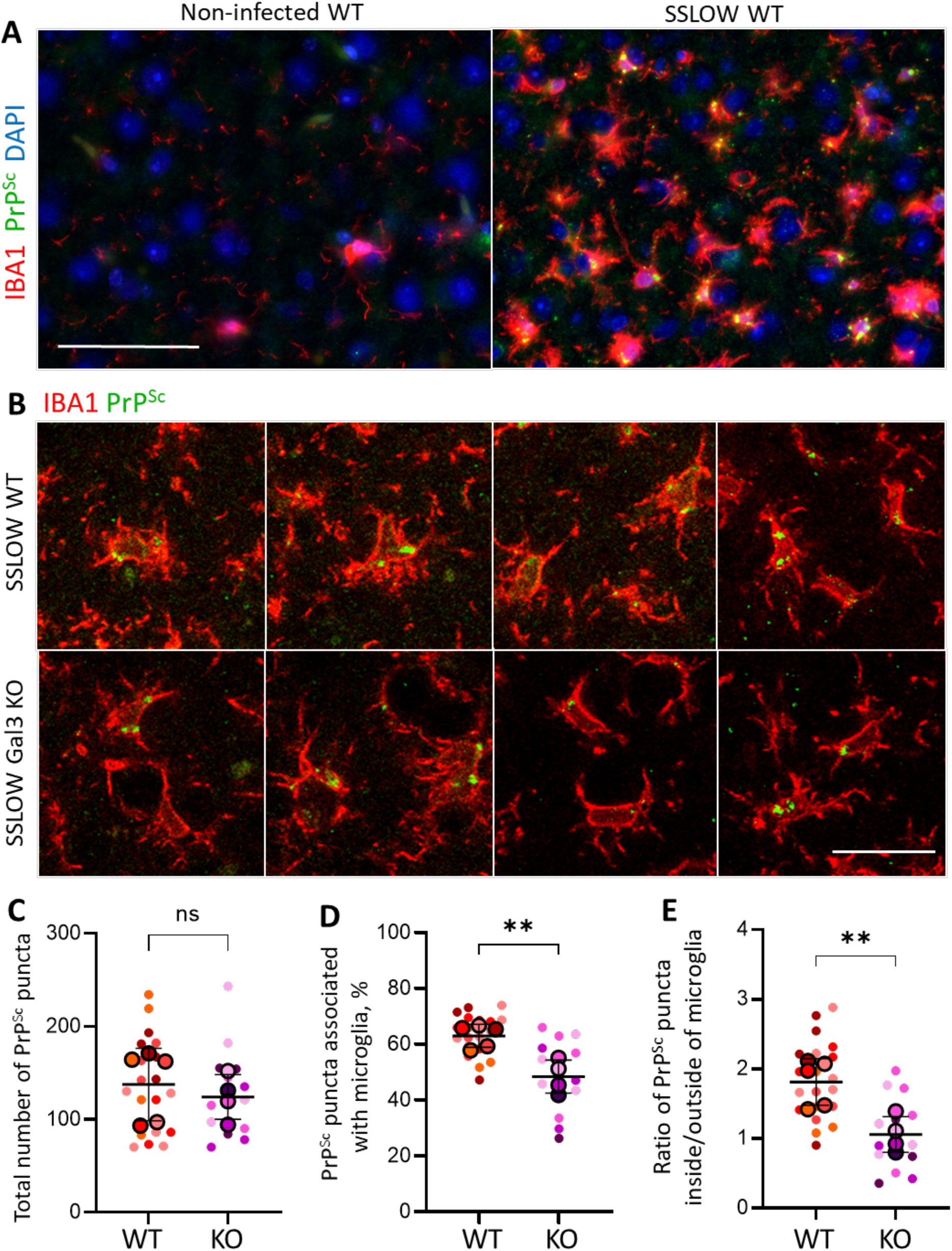
Analysis of neuronal envelopment by microglia. **A.** Confocal 3D reconstruction images of SSLOW-infected terminally ill Gal3 KO and WT mice showing examples of neuronal envelopment, in which microglia (IBA1, green) partially wrap around neuronal somata (NeuN, red). **B-E.** Quantification of neuronal density (**B**), percentage of neurons undergoing envelopment (**C**), microglial density (**D**), and percentage of enveloping microglia (**E**) in the cortex of SSLOW-infected terminally ill Gal3 KO and WT mice. N=5 animals per group, ns – not significant by unpaired Student’s t-test. Colors represent individual brains; dots represent individual fields of view; average values for each brain are shown as circles; means ± SD are marked by black lines. Scale bars 25 µm (top) and 10 µm (bottom).

### Gal3 deficiency reduces microglial uptake of PrP^Sc^

Because Gal3 is a component of the phagocytic machinery responsible for uptake of extracellular targets, including pathogens, cellular debris, and protein aggregates [56, 59], we next investigated whether Gal3 deficiency affects PrP^Sc^ uptake. Although microglia do not replicate prions *in vivo*, they acquire PrP^Sc^ positivity through phagocytic uptake [11, 12, 40]. We previously demonstrated that microglia internalize PrP^Sc^ beginning at early subclinical disease stages and accumulate substantial amounts of PrP^Sc^ within phagolysosomes by terminal disease [40].

Intracellular PrP^Sc^ puncta associated with microglia were readily detected in both SSLOW-infected Gal3 KO and WT mice (Fig. 5A). Quantification revealed no differences in combined extracellular and intracellular, microglia-associated PrP^Sc^ puncta between genotypes (Fig. 5B), consistent with Western blot analyses showing similar total PrP^Sc^ levels (Fig. 3B,C). However, the proportion of microglia-associated PrP^Sc^ puncta was markedly reduced in Gal3 KO mice relative to WT controls (Fig. 5D). Consequently, the ratio of intracellular to extracellular PrP^Sc^ was significantly lower in Gal3-deficient mice, indicating that Gal3 contributes to microglial phagocytic uptake of PrP^Sc^ (Fig. 5E).

**Figure 5.**
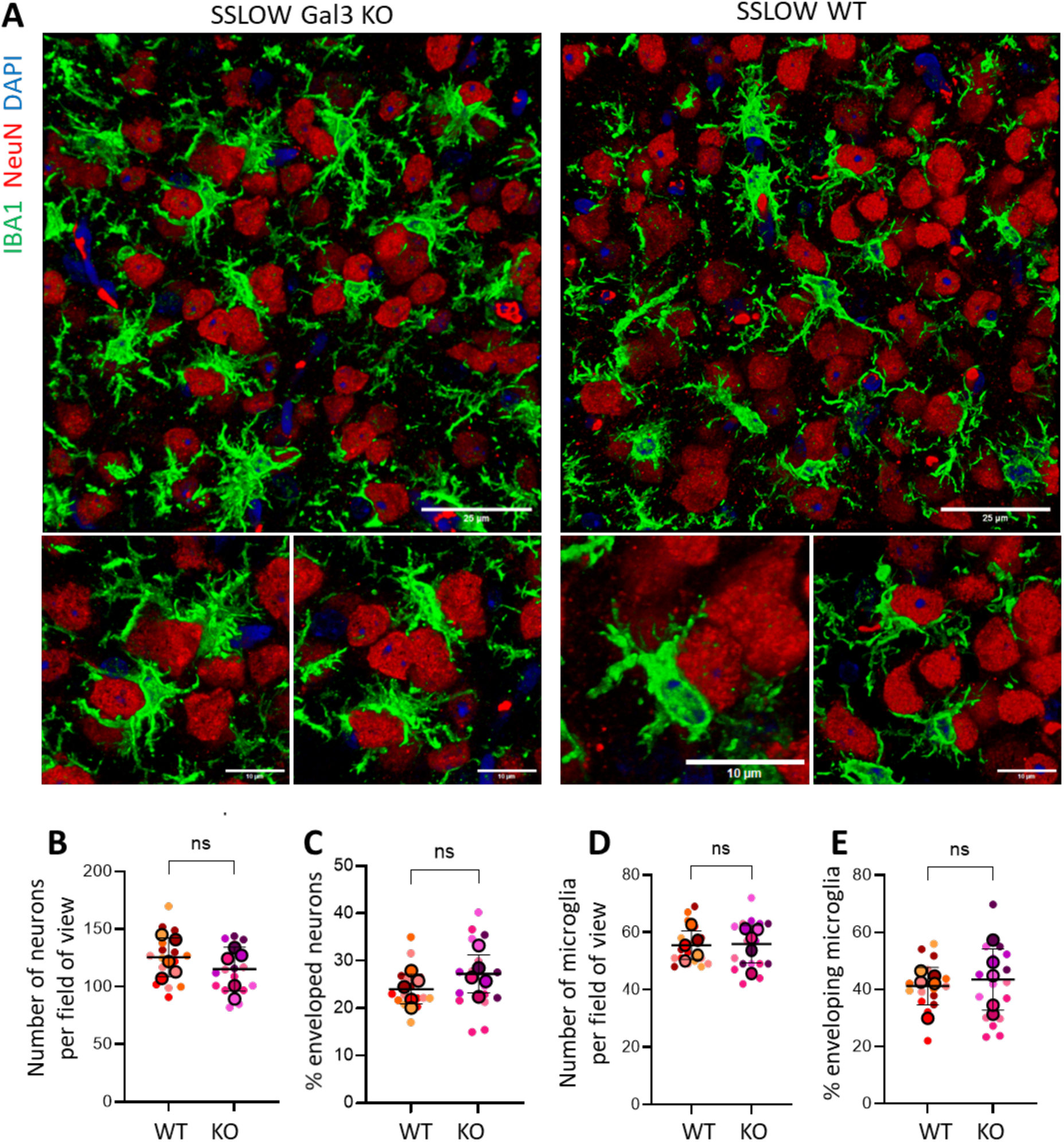
Analyses of PrP^Sc^ uptake by microglia. **A.** Representative immunofluorescence images from SSLOW-infected terminally ill C57Bl/6J mice and age-matched noninfected control mice stained for IBA1 and PrP^Sc^ (3D17 antibody). Scale bar 50 µm. **B**. Representative confocal microscopy images showing PrP^Sc^ (3D17, green) colocalized with microglia (IBA1, red) in SSLOW-infected terminally ill Gal3 KO and WT mice. Scale bar 20 µm. **C-E.** Quantification of the total number of PrP^Sc^ puncta (**C**), microglia-associated PrP^Sc^ puncta (**D**), and the ratio of PrP^Sc^ puncta associated with microglia per puncta not associated with microglia in SSLOW-infected terminally ill Gal3 KO and WT mice. Quantification was performed using integrated fluorescence densities of 3D17-positive puncta. N=5 animals per group, **p<0.01, ns – not significant by unpaired Student’s t-test. Colors represent individual brains; dots represent individual fields of view; average values for each brain are shown as circles; means ± SD are marked by black lines.

A recent study identified a subpopulation of microglia characterized by high expression of glycoprotein non-metastatic melanoma protein B (GPNMB) and transcriptional signatures associated with phagocytic activity [34]. This population was particularly enriched in the thalamus and spatially overlapped with *Lgals3*-expressing regions. To determine whether GPNMB and Gal3 mark the same microglial subpopulation, prion-infected C57BL/6J brains were co-immunostained for GPNMB, Gal3, and IBA1. Quantification within the thalamus, which exhibited the highest proportions of GPNMB ^+^ and Gal3^+^ cells, revealed partial overlap between these populations: approximately 20% of IBA1^+^ cells were Gal3^+^/ GPNMB ^-^, ∼10% of IBA1^+^ cells were Gal3^-^/ GPNMB ^+^, whereas 24% of cells were Gal3^+^/GPNMB ^+^ (Fig. S4A).

### Gal3 deficiency does not impair lysosomal functions

Accumulation of PrP^Sc^ within phagolysosomal compartments depends not only on uptake efficiency but also on degradation rates mediated by lysosomal activity. Protein aggregates, including α-synuclein, have previously been shown to disrupt lysosomal membranes and impair lysosomal function [60]. Gal3 has been implicated in sensing and promoting repair of damaged lysosomal membranes [29, 30, 61]. We therefore investigated whether Gal3 deficiency affects lysosomal function during prion disease.

Lysosomal activity was assessed by analyzing expression of LAMP1 and cathepsin D, both previously shown to be upregulated in microglia during prion disease [40, 62, 63]. Western blot analysis revealed no differences in expression of either protein between SSLOW-infected Gal3 KO and WT mice at either terminal or early clinical stages (Fig. 6A,B; Fig. S3A,C). In addition, the ratio of pro-cathepsin D (inactive precursor) to mature cathepsin D (active enzyme), a measure of lysosomal processing efficiency and autophagy-lysosome pathway activity [64], was unchanged between genotypes at both disease stages (Fig. 6B).

**Figure 6.**
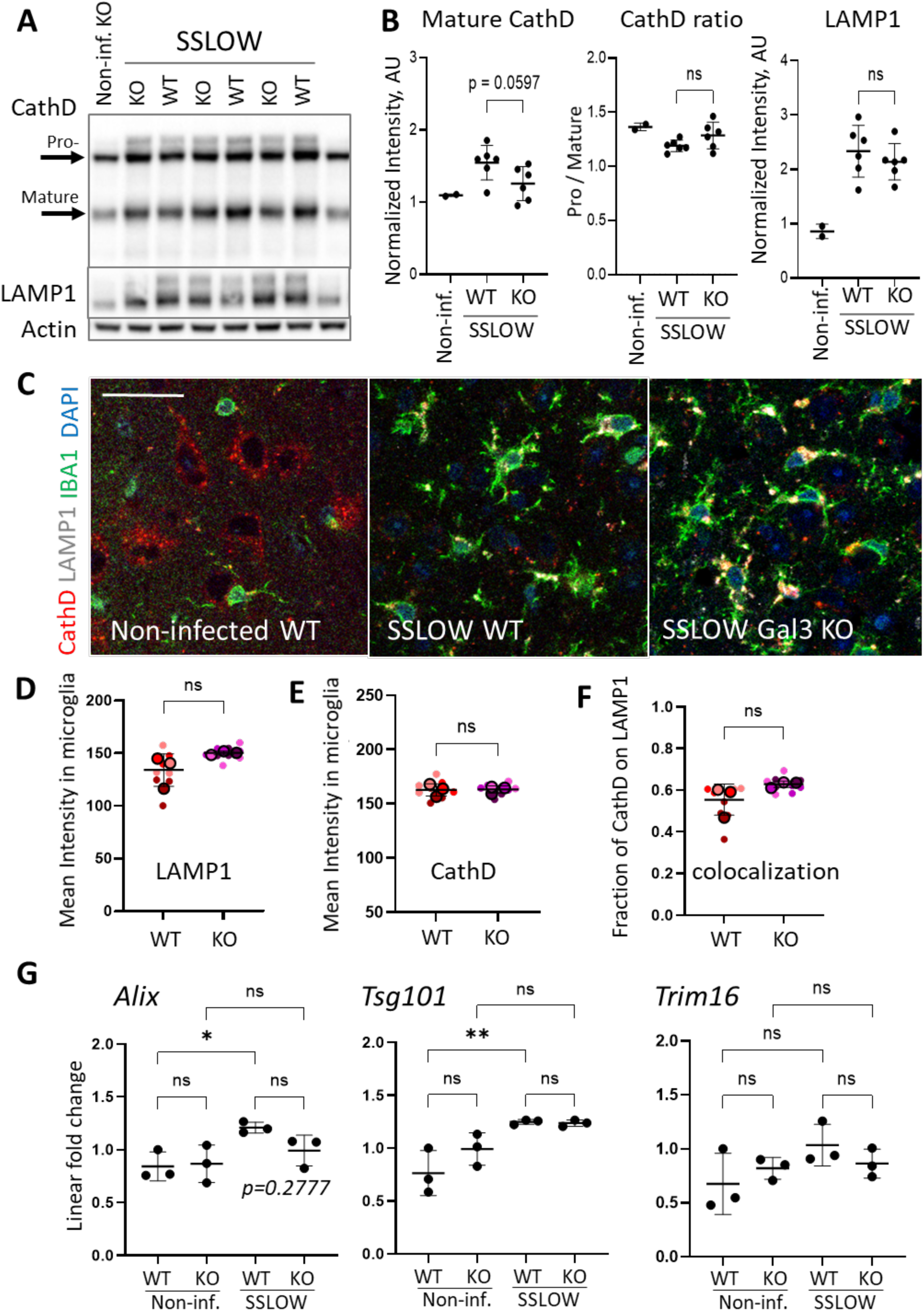
Analyses of lysosomal function in prion-infected Gal3 KO mice. **A.** Representative Western blot of whole-brain homogenates from SSLOW-infected terminally ill Gal3 KO and WT control mice. Arrows indicate pro-cathepsin D and mature cathepsin D heavy chain. **B**. Quantification of mature cathepsin D, the ratio of pro-cathepsin D to mature cathepsin D, and LAMP1 expression by Western blots. Protein levels were normalized to actin. N=6 animals per group, ns – non-significant, by unpaired Student’s t-test. Data from non-infected brains are shown as a reference (N=2). **C.** Representative confocal microscopy images showing colocalization of cathepsin D (CathD, red) with lysosomes (LAMP1, gray) within microglia (IBA1, green) in SSLOW-infected terminally ill Gal3 KO and WT mice, and age-matched non-infected WT mice. Scale bar 20 µm. **D-F.** Quantification of the mean fluorescence intensities of microglia-associated LAMP1 (**D**), CathD (**E**), and fraction of CathD colocalized with LAMP1 (**F**). N=3 animals per group. ns – not significant by unpaired Student’s t-test. Colors represent individual brains; dots represent individual fields of view; average values for each brain are shown as circles; means ± SD are marked by black lines. **G.** RT-qPCR analysis of gene expression in SSLOW-infected terminally ill Gal3 KO and WT mice and non-infected age-matched groups. N=3 animals per group, *p<0.05, **p<0.01, ns – non-significant, by ordinary one-way ANOVA with Tukey’s multiple comparison tests.

Immunofluorescence quantification of LAMP1 and cathepsin D within microglia confirmed that intracellular levels of these proteins were comparable between SSLOW-infected Gal3 KO and WT mice (Fig. 6C-E). Because lysosomal membrane rupture results in leakage of lysosomal contents and reduced colocalization of cathepsin D with LAMP1 [61], we next analyzed their colocalization within IBA1^+^ cells. Again, no genotype-dependent differences were observed (Fig. 6F). Finally, expression of genes involved in lysosomal function, sensing, and repair of damaged lysosomes (*Alix, Tsg101*, and *Trim16*) [29] was unchanged between genotypes (Fig. 6G). Together, these findings indicate that Gal3 deficiency does not measurably impair lysosomal activity or lysosomal membrane repair in prion-infected microglia. However, these results do not inform whether microglial lysosomes are damaged in prion diseases and whether their damage contributes to disease progression.

### Deficiency in microglial autophagy does not alter disease progression

Autophagy protects cells by removing damaged leaking lysosomes [65–67]. Therefore, next we investigated whether impairment of microglial autophagy influences prion disease progression. Beclin 1 acts as a central regulator of autophagy initiation promoting autophagosome formation around damaged organelles. To inhibit autophagy selectively in myeloid cells, we used homozygous Becn1^flox/flox^Lyz2^Cre/Cre^ mice (Becn1 cKO), in which Beclin 1 is deleted in myeloid lineages [42].

SSLOW-infected Becn1 cKO mice exhibited a modest reduction in incubation time to terminal disease relative to control Lyz2^Cre/Cre^ mice; however, this difference did not reach statistical significance (Fig. 7A). Both groups accumulated comparable levels of PrP^Sc^ as assessed by Western blotting (Fig. 7C,D). Confocal imaging demonstrated the presence of PrP^Sc^ aggregates within microglia in Becn1 cKO mice, similar to control and C57BL/6J mice (Fig. 7B,E). Quantification by immunofluorescence revealed a trend toward reduced microglia-associated PrP^Sc^ in Becn1 cKO mice, although the difference was not statistically significant (Fig. 7E).

**Figure 7.**
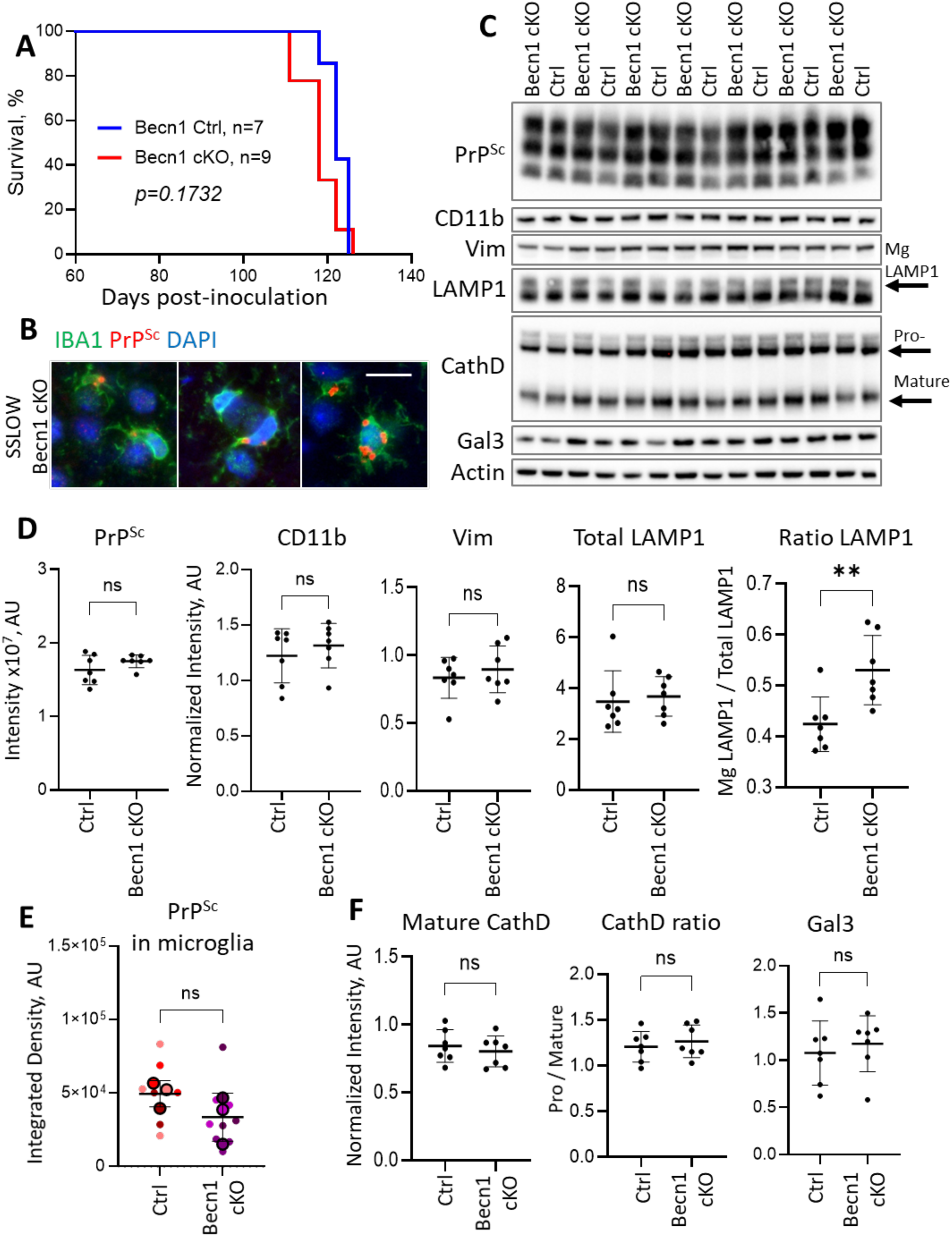
Analyses of disease progression in Becn1 cKO mice with impaired microglial autophagy. **A.** Survival curves of Becn1 cKO and control mice inoculated with SSLOW via i.c. route. Statistical analysis was performed using the Mantel-Cox log-rank test. **B.** Representative immunofluorescent images showing PrP^Sc^ puncta (SAF84, red) associated with microglia (IBA1, green) in SSLOW-infected terminally ill Becn1 cKO mice. Scale bar 10 µm. **C.** Representative Western blots of whole-brain homogenates from SSLOW-infected terminally ill Becn1 cKO and control mice. In the LAMP1 blot, the arrow marks the high-molecular-weight isoform attributed to microglia (Mg LAMP1). In the cathepsin D blot, arrows point at pro-cathepsin D and mature cathepsin D heavy chain. **D.** Quantification of PrP^Sc^, CD11b, Vim, total LAMP1, and the ratio of microglial LAMP1 to total LAMP1 in brains of SSLOW-infected Becn1 cKO and control mice by Western blot analysis. Protein levels were normalized to actin. N=7 animals per group, **p<0.01, ns – non-significant, by unpaired Student’s t-test. **E.** Quantification of integrated fluorescence densities of microglia-associated PrP^Sc^ in SSLOW-infected terminally ill Becn1 KO and control mice. N=3 animals per group. ns – not significant by unpaired Student’s t-test. Colors represent individual brains; dots represent individual fields of view; average values for each brain are shown as circles; means ± SD are marked by black lines. **F.** Quantification of mature cathepsin D, the ratio of pro-cathepsin D to mature cathepsin D, and Gal3 by Western blot analysis. Protein levels were normalized to actin. N=7 animals per group, ns – non-significant, by unpaired Student’s t-test.

No differences were detected between Becn1 cKO and control mice in expression of CD11b, Vim, total LAMP1, Gal3, cathepsin D, or the ratio of pro-cathepsin D to mature cathepsin D (Fig. 7C,D,F). Notably, microglial LAMP1, which is more heavily glycosylated than neuronal LAMP1 and can therefore be distinguished by Western blotting [68], was increased relative to total LAMP1 in Becn1 cKO mice, suggesting lysosomal accumulation within microglia (Fig. 7D), consistent with impaired lysophagy. Collectively, these findings indicate that inhibition of autophagy in microglia exerts only minor effects and does not substantially alter prion disease progression.

## Discussion

The current study demonstrates that Gal3 upregulation in prion disease becomes detectable at late preclinical stages and gradually increases with disease progression. Gal3 expression was observed in a subpopulation of reactive microglia and was most prominent in the thalamus, consistent with previous transcriptomic studies [34, 52]. The inverse correlation between cellular Gal3 levels and the expression of the homeostatic microglial markers P2Y12 and TMEM119 within individual cells further suggests that Gal3 is associated with an advanced reactive microglial phenotype.

A recent study proposed that the GPNMB^+^/Gal3^+^ microglial subpopulation is responsible for phagocytosis of apoptotic neurons and cellular debris [34]. In the present work, although both GPNMB and Gal3 were strongly upregulated in the thalamus, only partial overlap between GPNMB ^+^ and Gal3^+^ microglial populations was observed. These findings suggest that Gal3 is not essential for the function of GPNMB ^+^ microglia. Because the GPNMB ^+^ phenotype is thought to be induced by apoptotic or dead neurons [34], a process that primarily occurs during terminal disease stages, the temporal dynamics of Gal3 expression are particularly informative. Gal3 upregulation was detected substantially earlier than the onset of overt neuronal loss and instead closely followed PrP^Sc^ accumulation, which begins during preclinical stages [40]. These observations suggest that Gal3 expression is induced in response to PrP^Sc^ accumulation rather than neuronal death. Consistent with this interpretation, Gal3 deficiency did not alter close microglia–neuron body-to-body interactions, indicating that Gal3 is unlikely to participate in microglial docking to neurons.

In the current study, loss of Gal3 reduced PrP^Sc^ uptake by microglia and accelerated disease progression. Gal3 knockout animals displayed a significantly lower ratio of intracellular microglia-associated to extracellular PrP^Sc^ (Fig. 5), supporting a role for Gal3 in PrP^Sc^ internalization. Gal3 exhibits high affinity for uncapped β-galactose residues on branched glycans lacking terminal sialylation and can function as an opsonin that promotes phagocytic uptake [23–26]. Both PrP^C^ and PrP^Sc^ carry up to two branched N-linked glycans per molecule [69–72], with each glycan containing up to five negatively charged sialic acid residues [70–72]. In the mammalian brain, however, only ∼40% of N-linked glycan branches are sialylated, leaving a substantial proportion of uncapped β-galactose residues exposed [73]. Previously, we demonstrated that among the large diversity of PrP^C^ sialoglycoforms present in the brain, prion strains recruit PrP^C^ sialoglycoforms selectively [74–76], and that selectivity is dictated by N-glycan sialylation status [77]. Mouse-adapted strains, as well as PrP^Sc^ associated with distinct CJD subtypes, display a broad range of selectivity profiles. SSLOW is among the most selective strains, preferentially recruiting less sialylated PrP^C^ isoforms enriched in uncapped β-galactose residues [44]. Given that Gal3 mediates phagocytosis through recognition of uncapped β-galactose-containing glycans, the present findings provide experimental evidence that Gal3 contributes to the phagocytic machinery responsible for PrP^Sc^ uptake by microglia.

In addition to marked strain-specific differences [77, 78], PrP^Sc^ sialylation also varies across brain regions within the same strain [79]. Although these regional differences are modest, the relative ranking of PrP^Sc^ sialylation levels across brain regions is conserved across multiple strains [79]. Regardless of strain, thalamus-derived PrP^Sc^ consistently exhibits lower sialylation than PrP^Sc^ isolated from the cortex or hippocampus [79]. The observations in the current and previous studies that the thalamus displays the most pronounced neuroinflammatory response, including the strongest Gal3 upregulation [34, 48], together with the finding that thalamic PrP^Sc^ is the least sialylated [79], supports the hypothesis that microglia respond to exposed galactose residues on PrP^Sc^ glycans and that Gal3 contributes to PrP^Sc^ uptake.

Maintenance of lysosomal integrity is essential for phagocytic cells to internalize and process extracellular material. Previous studies in cultured cells demonstrated that Gal3 participates in sensing lysosomal membrane rupture and coordinating lysosomal repair and autophagic degradation [29]. We reasoned that accumulation of PrP^Sc^ within phagolysosomal compartments might compromise lysosomal integrity and thereby affect lysosomal function. However, we did not observe evidence of more severe lysosomal damage or impaired repair in prion-infected Gal3 knockout mice relative to wild-type controls. Expression levels of LAMP1 and cathepsin D, markers of lysosomal activity, were not altered by Gal3 deficiency. Moreover, colocalization of cathepsin D with LAMP1, a parameter indicative of lysosomal membrane rupture, was unchanged in the absence of Gal3. Expression of genes involved in lysosomal damage sensing and repair (*ALIX, TSG101*, and *Trim16*) was also comparable between prion-infected Gal3 knockout and wild-type animals. These findings suggest that, *in vivo*, Gal3 is not essential for maintaining lysosomal integrity during prion infection. One possible explanation is that the role of Gal3 in lysosomal repair, previously described in cultured cells [29], may differ under *in vivo* conditions, where additional compensatory mechanisms are likely to exist. Alternatively, lysosomal membrane repair pathways *in vivo* may be substantially more redundant than those operating in cultured systems.

Autophagy protects cells from toxicity by eliminating damaged and leaking lysosomes. To determine whether lysosomal damage influences disease progression, we examined prion disease in Becn1^flox/flox^Lyz2^Cre/Cre^ mice, in which autophagy is impaired through conditional deletion of Becn1 in microglia. An increased ratio of microglial LAMP1 to total LAMP1 signal illustrated that inhibition of microglial lysophagy was achieved in Becn1 cKO mice. Nevertheless, only a modest, statistically non-significant acceleration of disease progression was observed, suggesting that inhibition of autophagy in microglia has limited impact on disease progression. PrP^Sc^ accumulation within microglia begins during early preclinical stages [40]. In the absence of efficient degradation of PrP^Sc^, microglial proliferation may increase the overall capacity of the brain to sequester PrP^Sc^. However, despite this increase in sequestration capacity, PrP^Sc^ replication appears to become uncontrolled by the early clinical stage [40]. Thus, the ability of microglia to sequester PrP^Sc^ during subclinical stages likely influences disease trajectory and the timing of terminal disease. By contrast, autophagic clearance of damaged lysosomes may become relevant only at later stages, when individual microglial cells approach their storage capacity. At this point, preservation of phagolysosomal integrity may no longer substantially alter disease progression.

In summary, the present study demonstrates that Gal3 contributes to the uptake of PrP^Sc^ by microglia and that disruption of this pathway accelerates disease progression. Because substantial amounts of PrP^Sc^ still accumulated within microglia in Gal3 knockout animals, additional Gal3-independent mechanisms of PrP^Sc^ phagocytosis likely exist, providing functional redundancy for this critical microglial activity. Although Gal3 is upregulated only in a subset of microglia, its expression serves as a sensitive indicator of neuroinflammation across both disease stages and brain regions. Furthermore, Gal3 upregulation during disease progression is sensitive to alterations in microglial phenotype induced by deletion of microglia-associated genes.

## Acknowledgments

Financial support for this study was provided by National Institute of Health Grants R01 NS045585 and R01 NS129502 to IVB, and R01 NS115876 to MML.

## Conflict of Interest

The authors have no conflicts of interests to declare.

## Author Contribution

N.M. and I.V.B. conceived and designed the study; N.M., T.S., O.B., O.M., and N.P.P., performed experiments; K.M. managed mouse colony, performed animal procedures and scored the disease signs; N.M., T.S., and O.B. analyzed the data; M.M.L., provided animal model; I.V.B. and N.M. wrote the manuscript. All authors read, edited and approved the final manuscript.

## Data Availability Statement

The data that support the findings of this study are available upon reasonable request.

## Notes

### Competing Interest Statement

The authors have declared no competing interest.

