## Supplemental Figures for "Protective Role of Galectin-3 in Prion Disease Through Regulation of Microglial PrP^Sc^ Uptake"

**Figure S1.**

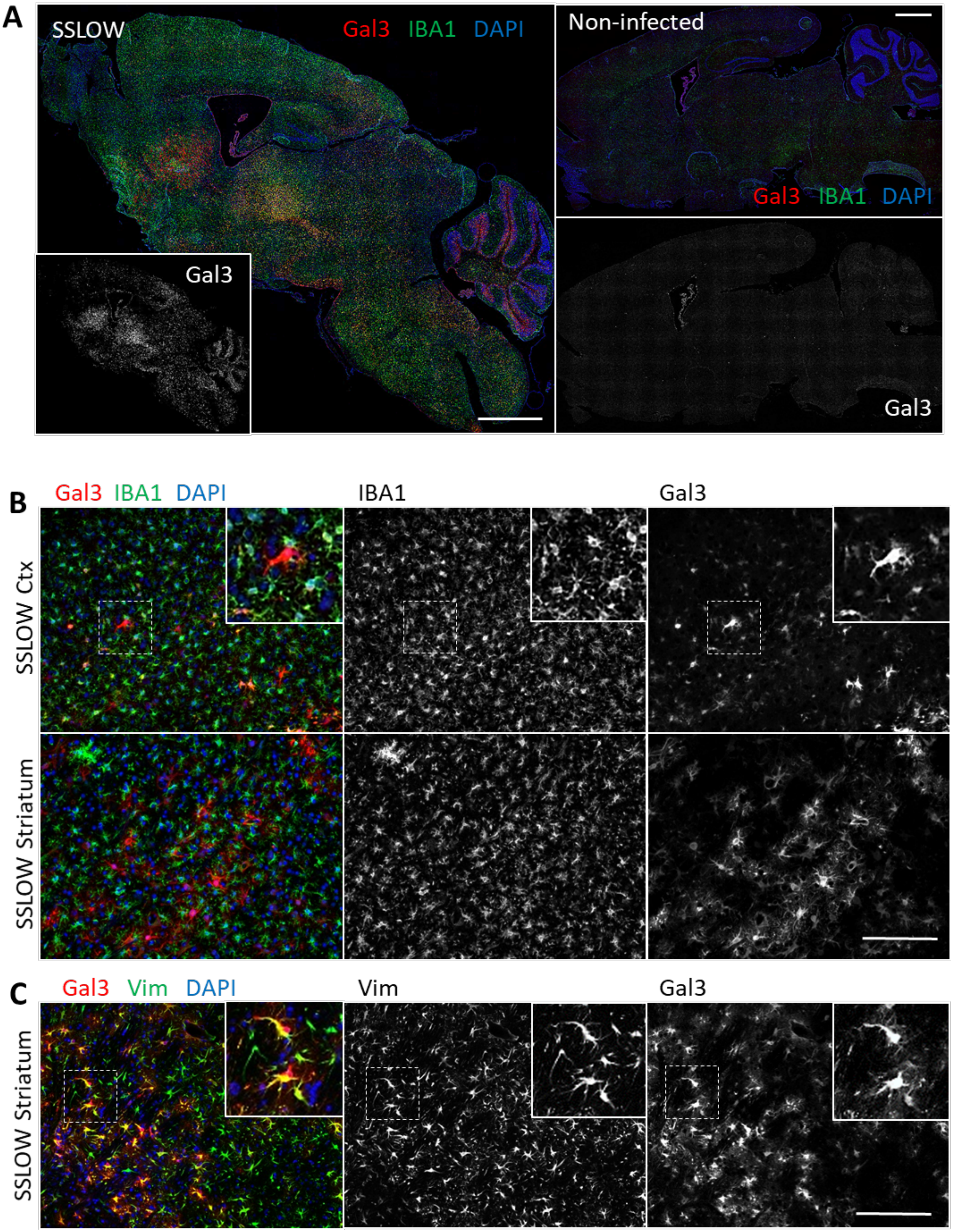

A
SSLOW
Gal3 IBA1 DAPI
Non-infected
Gal3 IBA1 DAPI
Gal3
Gal3
B
Gal3 IBA1 DAPI
IBA1
Gal3
SSLOW Ctx
SSLOW Striatum
C
Gal3 Vim DAPI
Vim
Gal3
SSLOW Striatum

**Figure S1. A.** Whole-brain merged mosaic images showing immunofluorescence staining for Gal3 and IBA1 in C57Bl/6J mice infected with SSLOW via i.p. and non-infected mice. Scale bar 1000 µm. **B.** Immunofluorescence staining of Gal3 in the cortex (Ctx) and striatum of SSLOW-infected C57Bl/6J mice. **C.** Co-immunofluorescence staining for Gal3 and Vim showing colocalization of Gal3 signal with Vim+ astrocytes in the striatum of SSLOW-infected C57Bl/6J mice. In **B** and **C**, high-magnification views of boxed areas are shown. Scale bar 100 µm in B and C.

**Figure S2.**

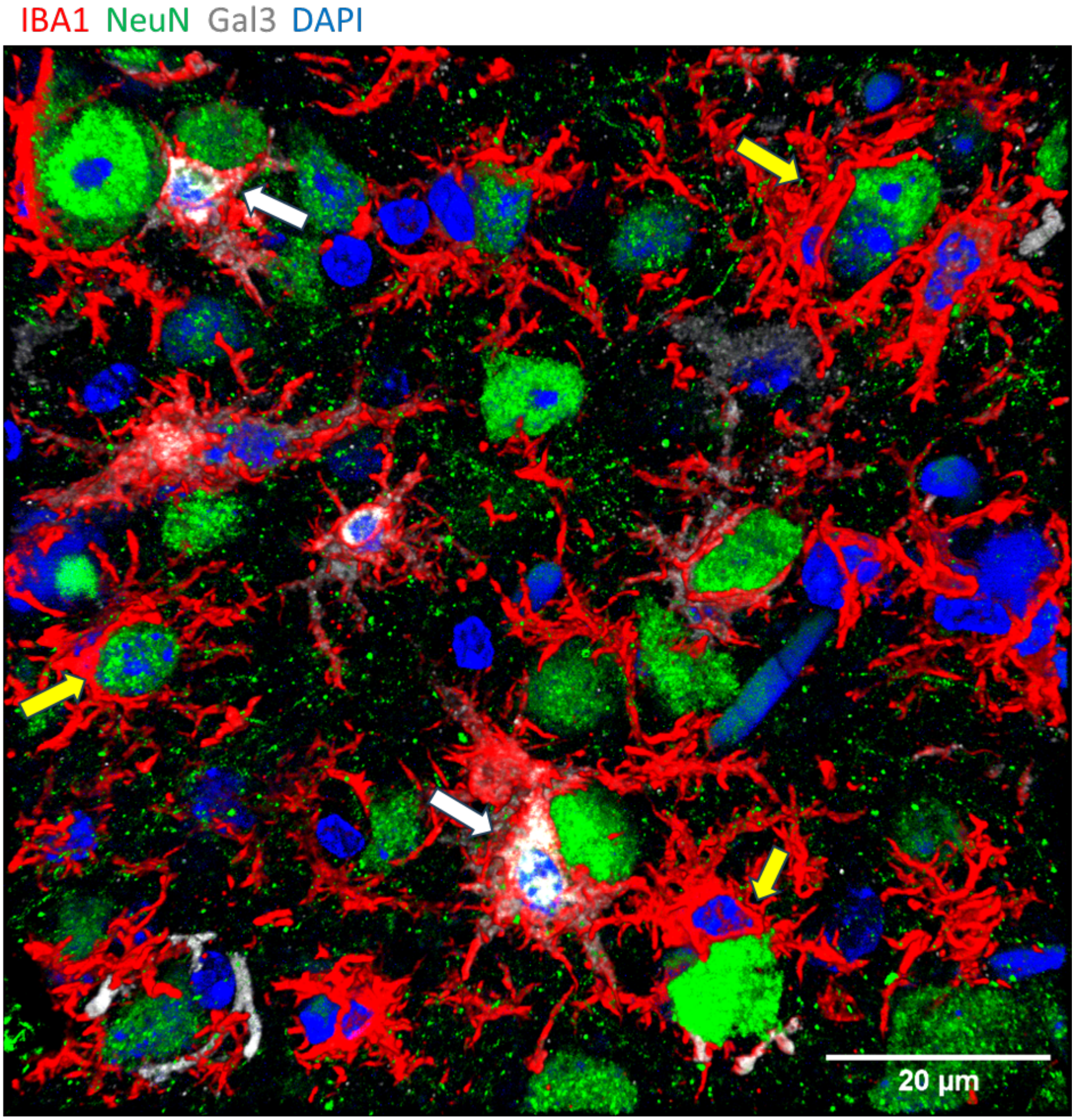


**Figure S2.** Confocal 3D reconstruction image showing Gal3-positive (white arrows) and Gal3-negative (yellow arrows) IBA1$^{+}$ microglia engaged in neuronal envelopment in the cortex of SSLOW-infected C57Bl/6J mice.

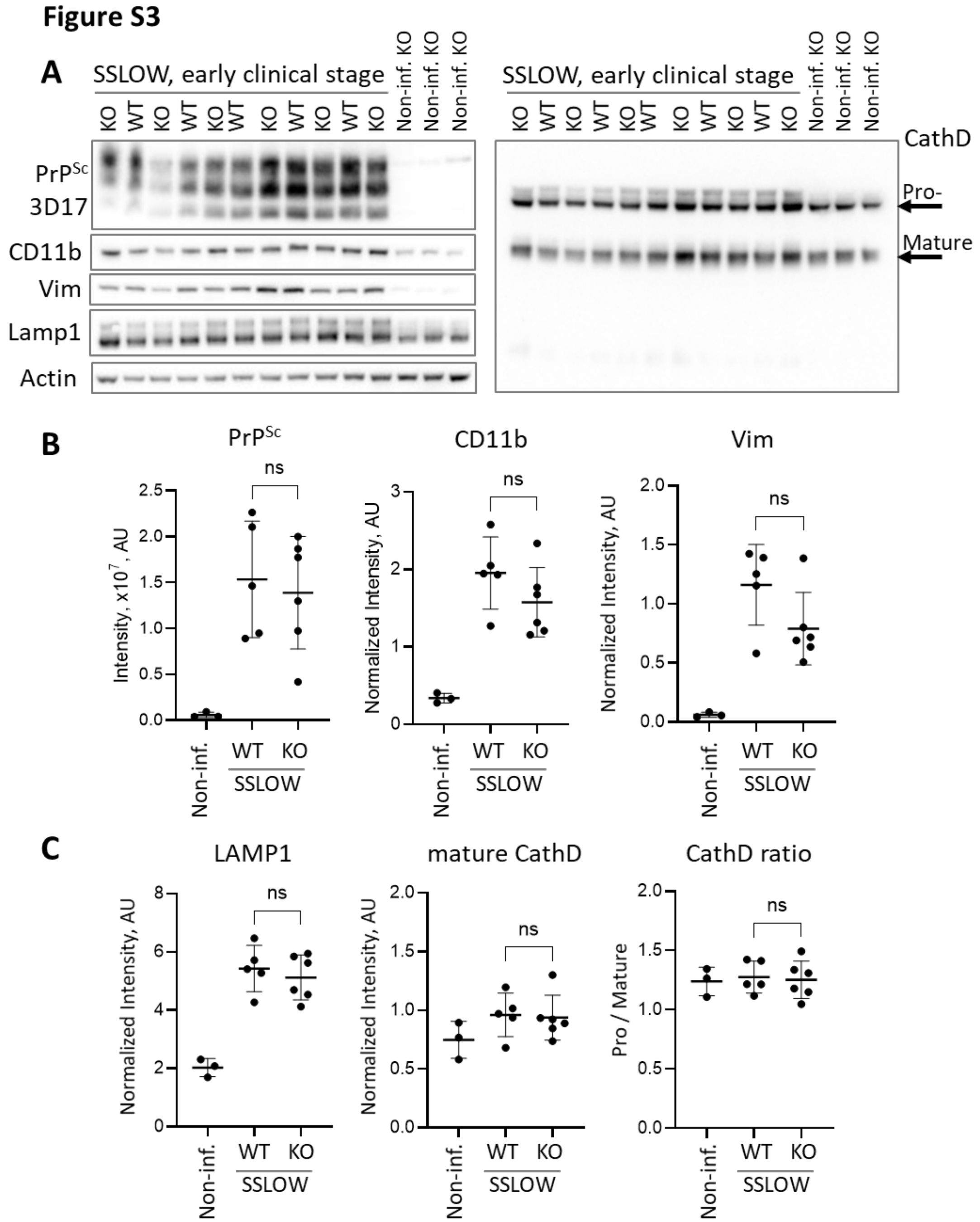


**Figure S3. Analysis of SSLOW-infected Gal3 KO mice at the early clinical stage.** Gal3 KO and WT control mice were infected with SSLOW via i.c. route and analyzed at 92-93 dpi. **A.** Representative Western blots of whole-brain homogenates from SSLOW-infected Gal3 KO and WT control mice, and non-infected Gal3 KO mice. **B.** Quantification of $PrP^{Sc}$, CD11b and Vim by Western blots. **C.** Quantification of LAMP1, mature cathepsin D and the ratio of pro-cathepsin D to mature cathepsin D by Western blots. In **B** and **C**, protein levels were normalized to actin. N=5-6 animals per group, ns – non-significant, by unpaired Student's t-test. Data for non-infected brains are shown for reference (N=3).

Figure S4

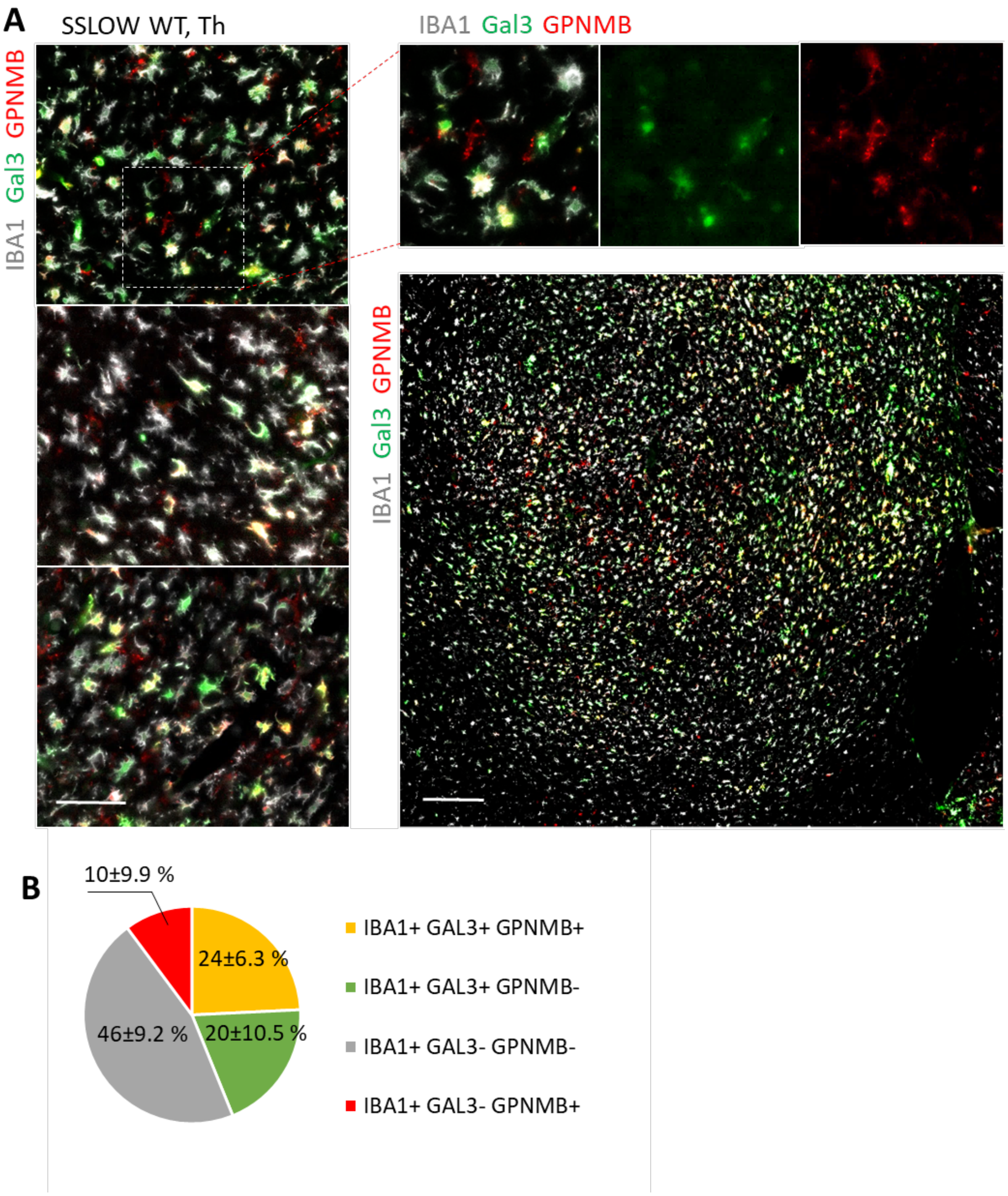


**Figure S4. Analysis of Gal3-positive and GPNMB-positive microglial subpopulations in prion-infected brain**. **A**. Representative immunofluorescence images of thalamic sections from SSLOW-infected terminally ill C57Bl/6J mice stained for IBA1, Gal3 and GPNMB. Scale bars 50 µm (left) and 200 µm (bottom right). **B**. Quantification of $IBA1^{+}/Gal3^{+}/GPNMB^{+}$, $IBA1^{+}/Gal3^{+}/GPNMB^{-}$, $IBA1^{+}/Gal3^{-}/GPNMB^{-}$, and $IBA1^{+}/Gal3^{-}/GPNMB^{+}$ microglial subpopulations within the thalamus. N=3 brains, with 111-124 cells per brain analyzed.
